# A Single-cell Spatiotemporal Manifold of Tissue Morphology and Dynamics

**DOI:** 10.1101/2025.10.22.683950

**Authors:** Erin Haus, Anthony Santella, Yichi Xu, Ruohan Ren, Dali Wang, Zhirong Bao

## Abstract

Complex tissues are now characterizable at single-cell resolution, but the spatial logic underlying tissue organization remains challenging to access without effective single-cell spatial descriptors. We present a learned single-cell manifold of cell positions over time, with which one can measure tissue morphology and trace dynamics. Learning co-processes pairs of cell point clouds sampled along time using a Transformer encoder with inter-sample attention, a strategy that promotes efficient joint spatiotemporal learning. The manifold shows desirable properties of a general descriptor, e.g. interpretable cell type clusters, preserved local distances, and a pseudo-time axis, and enables common but challenging spatial reasoning tasks such as annotation of anatomical landmarks at cellular resolution and detection of subtle, transient phenotypes in large screens. Our study demonstrates a widely applicable cell-based learning strategy and representation for studying tissue biology.

## Introduction

Complex tissues display diverse spatial configurations that evolve over development and disease progression. Imaging advances allow the observation of tissue dynamics at single-cell resolution in varied developmental and disease contexts [1-5]. In parallel, emerging spatial omics provides rich molecular information at single-cell resolution. However, it is still difficult to interrogate spatial relationship among cells in order to understand the patterns, schemes and underlying principles of tissue organization, as the spatial patterns are not necessarily evident in 3D. In spatial omics descriptors have been developed for regional cell type composition [6-8] helping define and measure neighborhood types, but the morphological relationships within and among neighborhoods largely remain to be explored. In contrast, manifolds in high-dimensional gene expression space have proven indispensable for tracing cell states and lineage differentiation as well as helping integrate multimodal omics data. A learned high dimensional space to represent tissue morphology could make cell organization and dynamics more accessible, and further enable integration of multimodal data.

We consider a cell-based learning approach, instead of pixel-based, to learn the desired space of tissue morphology. Cell position is a common data source for learning across modalities, and also a natural basis for potential multi-modal foundational models. A point cloud of cell positions is a general representation of tissue morphology at single-cell resolution and allows tissue measurement to be treated as the widely studied point cloud learning problem. Point descriptors of tissue, both traditional [9, 10] and learned [11, 12], have been used to align tissue samples, and can be further leveraged for systematic tissue morphology measurement. In the meantime, Transformer neural networks [13] have shown promise in representing spatiotemporal events including in point clouds [14]. Varied attentional strategies are being actively studied to address the computational and representational challenges of co-learning over space and time [15-18]. Measuring spatiotemporal events in tissues presents particular challenges, including large scale motion and variable timing of developmental events which result in variations of cell composition and context [19].

We present a learned manifold of single-cell embeddings created from cell point clouds to describe tissue morphology and dynamic changes. Specifically, we use a strategy we term Twin Attention for joint spatiotemporal learning, which trains a Transformer encoder to perform pairwise comparison of ‘twin’ samples drawn from a sliding local time window across individuals. The twinning strategy samples incremental developmental changes and biological variations to efficiently approximate global learning. We apply the method to *C. elegans* embryogenesis captured as dynamic point clouds of nuclear centroids that trace the proliferating cell lineage (Fig. 1a & S1a,b). The resulting manifold sorts distinct cell types, reflects neighborhood structure and global shape, and contains a pseudo-time axis consistent with development. We further demonstrate the power of this representation in two spatial reasoning tasks that are time consuming and have been difficult to automate: annotation of anatomical landmarks, which at the single-cell level provides cell type assignments, and phenotype detection. We suggest this learned manifold provides a general and interpretable representation of tissue morphology and dynamics valuable in probing their underlying spatial logic.

**Figure 1.**
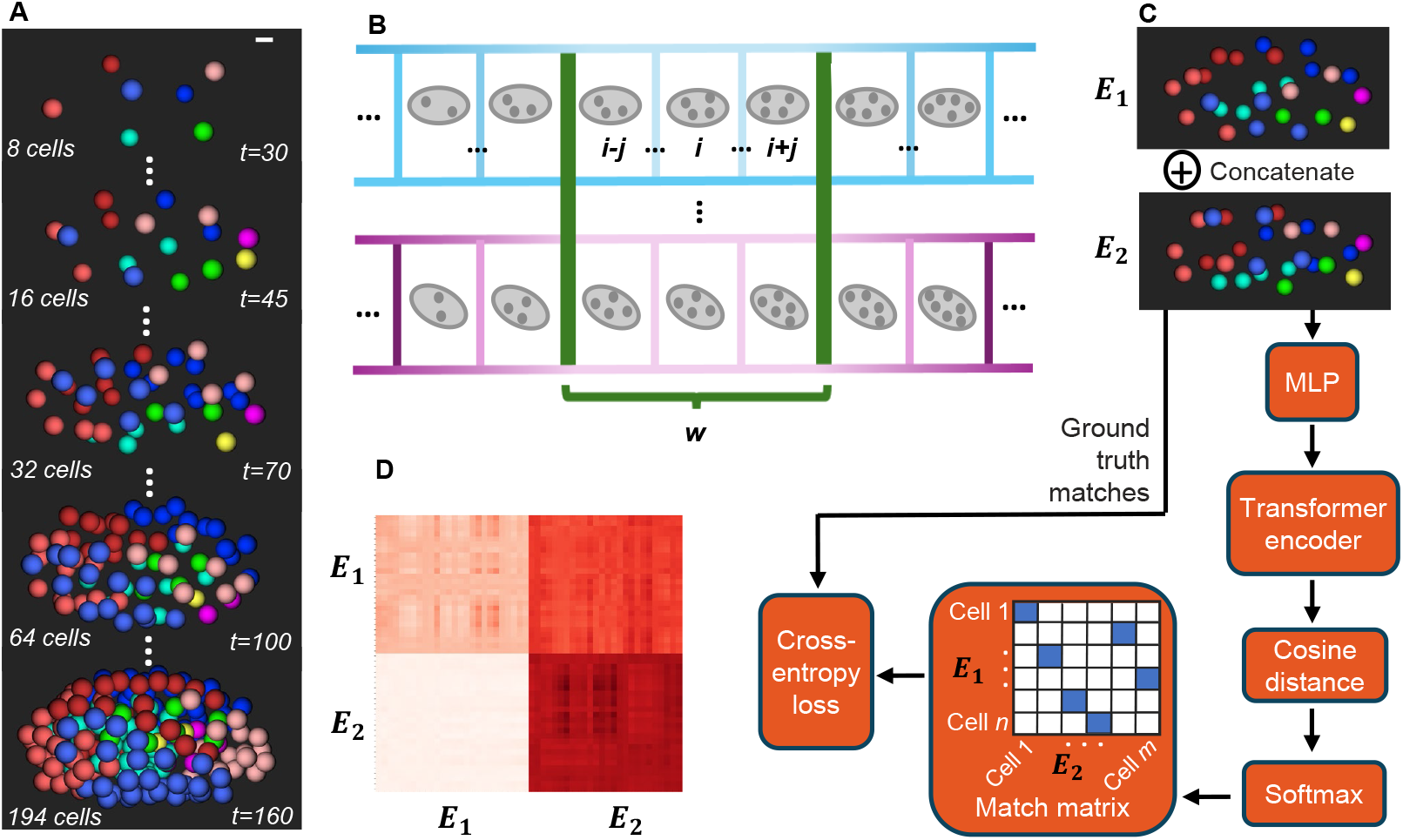
Learning a single-cell developmental manifold of tissue morphogenesis. A. Representative time points from a time series of early *C. elegans* embryogenesis. Spheres represent nuclei, colored by their lineages (red:ABa, blue:ABp, pink:C, magenta:D, green:E, cyan:MS, yellow:P). Time of development is indicated by *t* (minutes). Scale bar 5μm. For additional information on *C. elegans* embryogenesis, see Figure S1A-B. B. Sliding time window-based organization of training data for a local joint spatiotemporal learning strategy. A sliding time window (denoted ‘*w’)* is employed to define pools of embryonic point clouds. Pairs of point clouds are randomly sampled from each pool for training. Time series of embryogenesis are shown schematically in gray, with *i*, and *j* denoting time. C. Twin-attention for joint spatial and temporal learning. A pair of embryonic point clouds (***E***_**1**_ and ***E***_**2**_) are concatenated and presented to a Transformer encoder, which is trained to maximize the similarity between corresponding cells. The example embryos ***E***_**1**_ and ***E***_**2**_ are within the 2\**j*-cell-sized window around the *i*=26-cell stage, containing exactly *n* and *m* cells, respectively. For training performance, see Figure S1D. D. Inter and intra-sample attention map of the first layer of the Transformer encoder after training, for two embryonic point clouds (***E***_**1**_ and ***E***_**2**_) around the 26-cell stage. Intensity is the attention of cells along rows to those in columns averaged across the multiple-attention heads. Attention in additional layers of the Transformer encoder are provided in Figure S1E.

## Results

### Strategy and Base Training

A pair of cell point clouds, or ‘twins’, sampled from a local time window and across individuals (Fig. 1b) can be viewed as a mixture of invariant cellular configurations and developmental changes, plus biological variations specific to each sample. Developmental changes include cell motion and cell division. Variations include cell position and timing of developmental events known as heterochrony. Heterochrony of cell divisions, in particular, can create many possible combinations of cells and a highly variable context for learning (Fig. S1c,d). To facilitate comparison, we use an approach [11] that trains a Transformer encoder to match corresponding cells, based purely on x,y,z position, by forcing matched points to have similar embeddings (Fig. 1c). Concatenation of the pair enables both intra- and inter-sample attention (Fig. 1d). Timing difference in the pair leads to joint spatiotemporal learning, and the sliding window approximate global learning. Subtle timing differences and heterochrony are ubiquitous in staged samples regardless of species and anatomical level; the ability to create consistent representations of structures with inconsistent context would have broad implications in terms of spatiotemporal learning of tissue morphology.

We first perform base training on simulated data. A thousand embryos are simulated through the 194-cell stage, using an agent-based simulator that draws from measured distributions of cell cycle length and average migration paths [22]. The simulated data are further augmented 10x with random rotations (see Methods for training details). The trained base model successfully identifies correct matches on held-out data given only cell positions; accuracies are typically above 99% on matchable cells (present in both point clouds), even during periods where heterochrony of cell divisions leads to 30-40% of the cells not being shared between a pair of point clouds (Fig. S1d). This success is despite a challenging learning task where data is limited both in raw quantity and in sampling the combinatorial variation of cell composition caused by divisions. Notably, examination of the attention map suggests that the Transformer successfully manages the inter- and intra-sample attention (Fig. 1d & S1e).

### Spatiotemporal Properties of the Manifold

Twin Attention learning results in a manifold of single-cell embeddings over space and time. Firstly, training has driven distinct and robust embeddings for individual cell types. When embeddings are visualized in 2D with t-SNE (Fig. 2a), they are clustered by cell type despite biological variability (e.g., highly variable cell composition), embryo orientation, and cells existing across a range of developmental stages. Secondly, cells that are closer in real 3D space are closer in embedding space. Examining a narrower time window when a cell divides (Fig. 2b), the daughter cells are closest to the mother cell cluster in embedding space as they are in real space, even though mother/daughter pairs are not considered matches during training. Thirdly, there are clear temporal trajectories in the manifold. Embeddings for a lineage branch, i.e. a single cell over multiple rounds of division (Fig. S1a), change gradually over time (Fig. 2c), sweeping out a curve in the embedding space over generations. The global consistency in space and time suggests that the Twin Attention strategy achieves the expected joint spatiotemporal learning.

**Figure 2.**
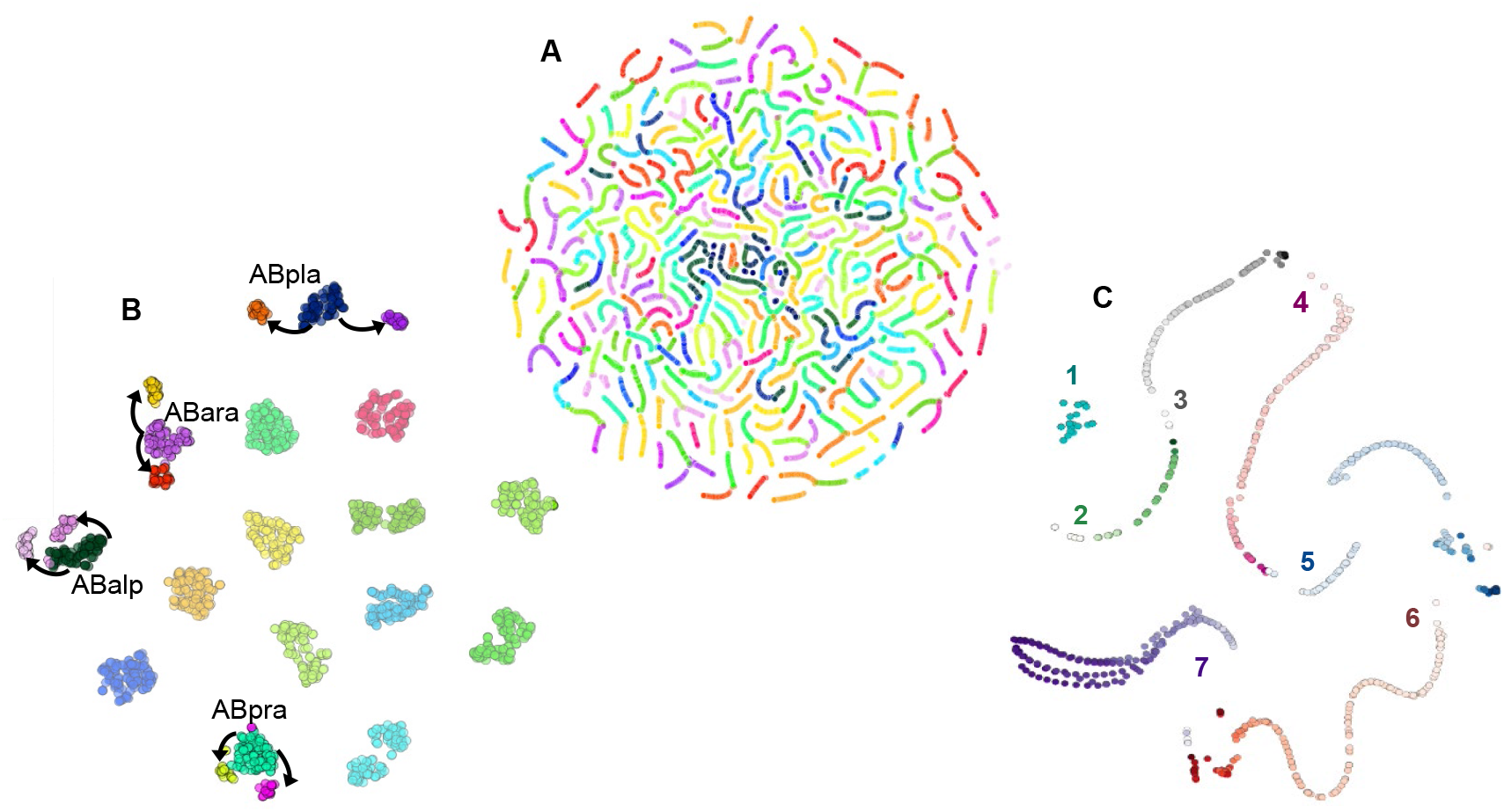
The learned single-cell manifold of tissue morphogenesis. A. Distinct and robust embeddings for cells. t-SNE of embeddings for all cells throughout development (4-to 194-cell stages, n=20 embryos, perplexity=50, random color per cell type). B. Vicinity in real and embedding space. t-SNE of embeddings for cells at a single stage (16-cell stage, n=20 embryos, perplexity=20, random color per cell type). Sub-clusters of mother and daughter cells are labeled with mother cell name and arrows toward daughter cell clusters. Different cells have divided in different embryos, highlighting heterochrony. C. Temporal trajectory in the manifold. t-SNE of embeddings for cells along a single lineage path (ABa-ABalaaaal, as colored in red in Figure S1A) (n=3 embryos, perplexity=20, random color per cell type). Color darkens forward in time per cell type. Numbers correspond to cell type/generations of cells along the lineage, with 1 denoting ABa and 7 denoting ABalaaaal.

We further examine the interpretability of the embedding in terms of space, time and cell-cell relationships. Spatially, we find that distance between embeddings preserves local distances in real space. At close distances a linear relationship exists between pairwise cell distances in real and embedding spaces, while this plateaus at greater distances (Fig. 3a). Such a relationship is preserved over a range of developmental stages (Fig. S2a). We use a sigmoid fit to approximate this relationship and consider the point at which the fit achieves 90% of its asymptotic value a point of transition. Over developmental stages, this transition point falls within a narrow range of values between 2 and 4 mean nearest neighbor distances. We also find that embedding distances over time reflect invariance to local density. While distance in real space decreases as cells divide and become smaller and denser, distance in embedding space remains constant (Fig. 3b). This relationship is consistent over different lineage branches (Fig. S2b). Together these observations suggest the manifold closely reflects local neighborhood structure.

**Figure 3.**
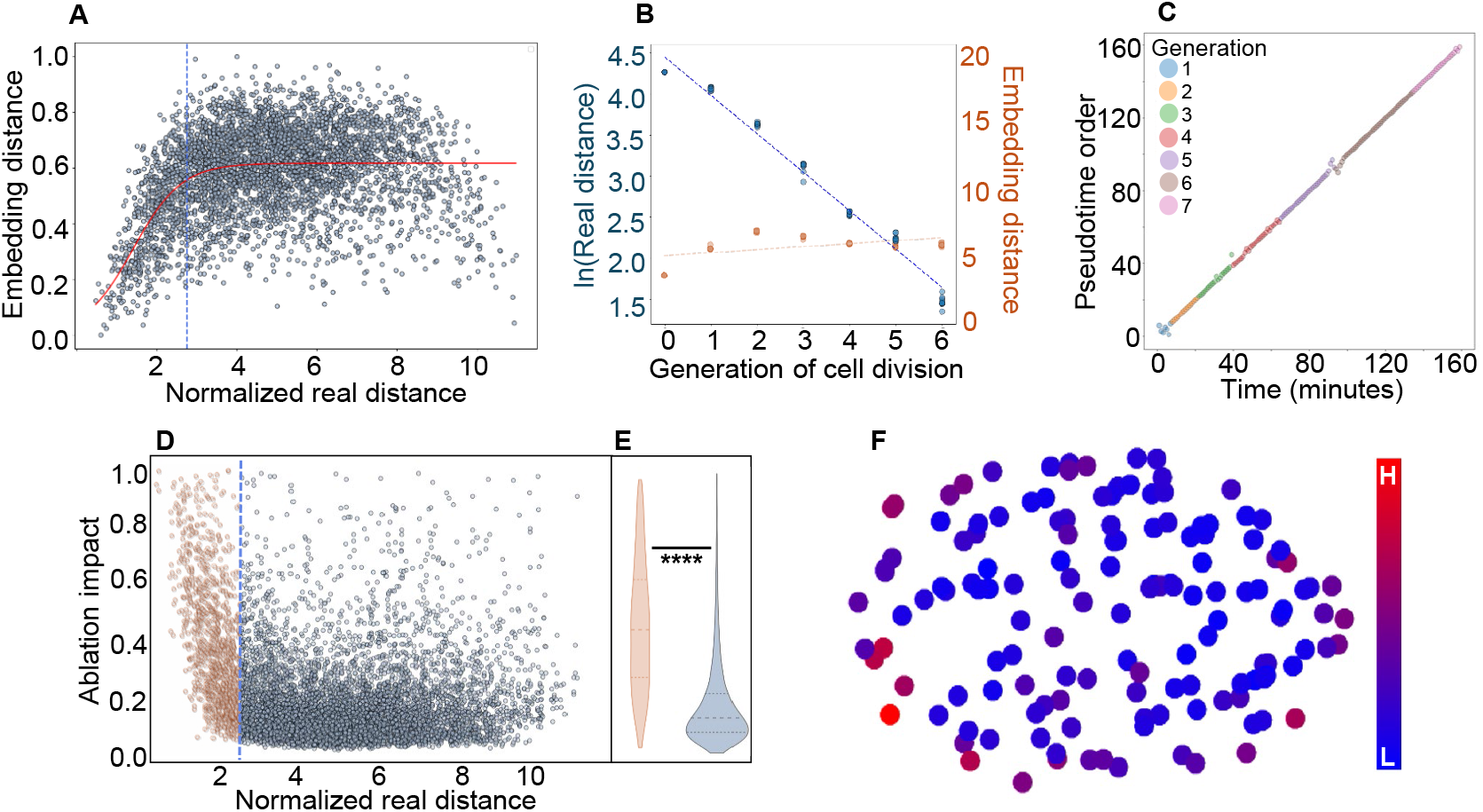
Characterization of the developmental manifold. A. Embedding distance reflects local distance in real space. Pairwise cell distance in real 3D and embedding space, shown for a 90-cell stage embryo. Red line is the fit with a sigmoid function. The transition point in the relationship, defined as the x-value at which the sigmoid function reaches 90% of its asymptotic value (blue line), indicates where the direct relationship between distances across the spaces diminishes. Distance in real space is normalized by the average nearest neighbor distance at this stage, reflecting layers of neighbors. Distance in embedding space is normalized by the maximum value. For additional developmental stages, see Figure S2A. B. Scaling with density. Distance between sibling cells in real and embedding space. The median pairwise distance in real (blue) and embedding (orange) spaces over generations along a sister-lineage path (MSppp(a/p) and sibling (n=10 embryos). Distance in real space is plotted in natural log scale. Dashed lines are linear fits. For summary of all lineage paths, see Figure S2A. C. Pseudotime axis. A single cell’s temporal trajectory through embedding space. Pseudotime is computed from the embeddings of cells along a single lineage path (ABa-ABalaapaa) and is defined here as the order of the points along a 1D diffusion map reduction of the embeddings. For a summary of all lineage paths, see Figure S2C. D. Local influence of in silico cell ablations. Influence of cells at varying distances on embeddings, shown for a 90-cell stage embryo. Pairwise cell distance in real space versus the change in embedding position before and after a cell’s removal (“ablation impact”, or Euclidean distance in embedding space before and after ablation). Distance in real space is normalized as in (A). Impact normalized by max value. Orange indicates nearby neighbor pairs, distinguished from further neighbors (gray) by the transition point as described in (A). For additional developmental stages, see Figure S2F. E. A summary of ablation impact in (D). Violin plots of near and distant effect of ablations (grouped by values less or greater than the transition point in (A)), with mean and quartiles marked by dotted lines. Stars denote statistical significance of the difference (Welch’s t-test, p=5.27e-120). For additional developmental stages, see Figure S2G. F. Some cells show global impact. Summary of per cell ablation impact. Summed impact of each cell on all other cells when computationally ablated, 140-cell stage colored by impact. For additional developmental stages, see Figure S2I.

To further examine the temporal trajectory in embedding space (Fig. 2c), we use a diffusion map [23] to reduce the manifold of a lineage branch (Fig. S1a) to one dimension as an approximation of the pseudo-time axis. Ordering in the diffusion map shows a near perfect monotonic relationship with real time (Fig. 3c). This relationship holds for different lineage branches (Fig. S2c). However, distance along the 1d diffusion map is not entirely linear with real time (Fig. S2d), which may be due to the oversimplification of dimension reduction.

To investigate cell-cell relationships, we conduct *in silico* cell ablations to quantify how cells contribute to each other’s embedding. In ablation, each cell is removed in turn and embeddings for all other cells with and without that cell are compared (Fig. S2e). Ablated cells have a larger impact on their neighbors (Fig. 3d, S2f). Nearby populations show consistently higher impact across developmental stages (Fig. 3e, S2g), where nearby is defined as closer than the transition points above (Fig. 3a, S2a). Overall impact of an ablation, which sums the change of every cell, is variable (Fig. S2e,h). A few cells impact most embeddings, while the majority have limited impact. The most influential cells appear to be concentrated at spatial extrema of the point cloud, particularly along the longest tissue axis, i.e., the anterior and posterior poles (Fig. 3f, S2i). Neighbors remain more impacted whether total impact is high or low (Fig. S2h). Thus, the manifold reflects both local relationships and the global distribution of cells.

Based on the analyses above, we conclude that the Twin Attention strategy successfully learns an interpretable single-cell manifold of tissue morphogenesis.

### Applications: Overview and Fine-Tuning

To examine the power of the manifold in studying tissue morphology and dynamic changes, we explore two common and challenging spatial reasoning applications, namely annotation of anatomical landmarks and phenotype detection, at the cellular level.

Prior to the applications, we fine-tune the base model on real data, which are available but limited. We use two datasets, the first 49 embryos from a previous study up to the same 194-cell stage as in the simulation experiment (dataset 1) [1, 24], the other 233 embryos up to the 520-cell stage generated in another lab with a similar imaging protocol (dataset 2) [25]. Training starts with base model weights but is otherwise identical (see Methods). Fine-tuning is successful in that performance of pairwise cell matching is above 80% for matchable cells including the > 500-cell stage (Fig. S3a). The manifold of the real data (Fig. 4a) shows distinct clusters for cell types as for the simulated data, albeit noisier (Fig. 2a). We note that after fine-tuning up to the 520-cell stage, over a thousand cell types are successfully embedded and distinguishable. These results demonstrate generalizability of the base model over different developmental stages and experimental setups.

**Figure 4.**
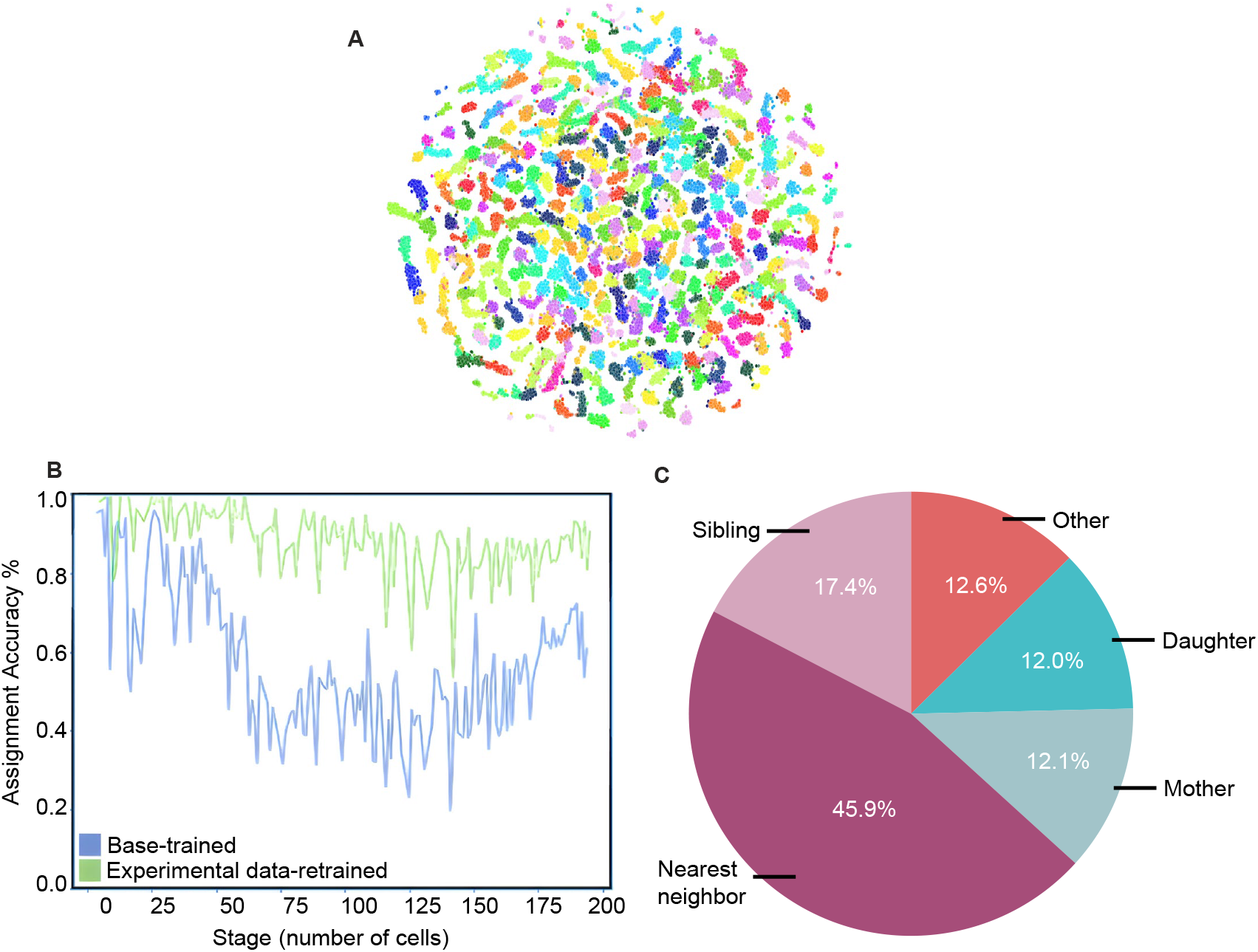
Fine-tuning of base model for cell type assignment. A. Manifold of embeddings from fine tuned model (dataset 1). Fine tuning for real data generates distinct and robust embeddings for cells. t-SNE of experimental embryos that make up the fine-tuning and kNN basis dataset (4-to 194-cell stages, n=39 embryos, random color per cell type, perplexity=50). B. Cell type assignment accuracy from kNN classification over stages (dataset 1, n=10 held-out embryos). Results are shown for manifold resulting from training on purely simulated data and that fine-tuned using experimental data. C. Error classes of identity predictions on the held-out embryos (dataset 1, n = 10 embryos at the 4-to 194-cell stages). Misassignments can be separated into three major classes: 1) close neighbors (purple), a special class of this case being sister cells, 2) a cell’s own mother or daughter (blue), and 3) confusions (red) that arise from any other scenario.

### Application 1: Position-Based Annotation of Anatomical Landmarks

Landmark annotation is a common task in image analysis ranging from cell level brain atlas generation to whole body organ segmentation. At the cellular level, the task becomes assigning cell type tags to individual cells. Because of the challenge of assembling appropriately specific marker panels, as well as the technical difficulties and costs of imaging multiplex markers, assignment based on cell positions is highly desirable. In *C. elegans*, the cell type tags are the known lineage identities [26]. The most reliable approach is live imaging and direct lineage tracing, which is not only technically challenging to do but also not amenable for fixed samples.

We use the learned embeddings to assign lineage identities based purely on cell positions. Specifically, we use embeddings from multiple embryos as the reference distribution, and treat cell type assignment as a k-nearest neighbors (KNN) classification task. For dataset 1, the average KNN accuracy (k=30) per point cloud over all stages is 88.2% and a significant improvement compared to using the base model trained on simulated data (Fig. 4b). For dataset 2, the average accuracy per point cloud over all stages is 93.0% up to the 520-cell stage (Fig. S3b), likely representing improvement from a larger training set and reference distribution. Notably, variation of cell composition affects the performance. At stages with no or few variable cells, the accuracy is about 95% or higher. This performance, despite a simple kNN classification, is a significant improvement on cell identification in *C. elegans*, including our own prior work in late stage embryos using a more traditional alignment method (71-78%) [19] and recent efforts to annotate about 50 to 200 labeled neurons in adult whole brain images based on positions (< 75%) [11, 27].

Assignment errors fall into three categories (Fig. 4c). The first and largest is confusion with spatial nearest neighbors, including the special case of sibling cells which are typically the closest neighbor. These combined constitute 63.3% of the errors. The second largest class is inter-generational confusion between mother and daughter cells, which are temporal nearest neighbors. These cases account for 24.1% of the errors. The third class is confusion with more spatially or temporally distant cells, constituting 12.3% of the errors. Confusion with the nearest spatial or temporal neighbors is consistent with the interpretable properties of the manifold, and the limited number of nonlocal identity confusions further suggests the manifold is largely non-self-intersecting.

Overall, strong performance demonstrates that the embeddings can provide accurate and robust landmark annotation. We note that the cell type assignment task here is rather challenging. To casual inspection, cells appear to be uniformly distributed in the embryo, so the spatial patterns are subtle. Furthermore, dozens to hundreds of cells are present in a point cloud with extreme variability of cell composition (up to 40% cells being variable), which present a large and complex combinatorial problem. Therefore, the learned embeddings must be richly descriptive and robust to contextual variability.

### Application 2: Detection of Subtle and Transient Phenotypes

Our second application seeks systematic and automated measurement of morphogenesis phenotypes in large-scale screens. We aim to detect subtle and transient phenotypes. Such phenotypes are of interest as they may reveal alternative developmental trajectories to the same end state, or error correction mechanisms for developmental robustness, but have been difficult to find as traditional screens typically rely on gross terminal phenotypes. For full automation, we further challenge our method by using point clouds with cell detection errors, which is typical of automated cell detection.

Specifically, we examine a set of RNAi treated embryos collected, but not analyzed, from a prior study [28]. This dataset contains 703 embryos which survived treatment with RNAi for 144 different embryonic lethal genes (see Supplementary File 1 for list), likely due to RNAi inefficacy. Manual curation of the automated lineaging results [29, 30] on these embryos would take over 500 person hours, an overwhelming effort given the likeliness of only finding wild-type (WT) behavior. We take the automated cell detection results corrupted by detection errors, and without the cell type/lineage identity assignments.

Phenotype detection based on the WT-trained embedding is a challenging out-of-distribution test of our method, a classic test for overlearning. We reason that with subtle phenotypes, cell embeddings of the RNAi treated embryos would be close to, but distinguishable from the corresponding WT cells. To test this notion, we assign cell types for the RNAi treated embryos as above, then measure distances to the corresponding WT cells. We find that distance from the manifold is greater for RNAi treated embryos than for an independent set of WT embryos with detection errors (Fig. 5a). To evaluate the cell assignment, we curated 32 of the RNAi treated embryos to establish correct cell type assignments. While accuracy for identifying individual cell types was moderate, the accuracy in assigning cells to founder lineages is above 80% (Fig. S4.1a). We conclude that our notion is correct, and that phenotypes can be detected and localized to specific tissues using the WT manifold albeit at a coarser level of lineage groups.

**Figure 5.**
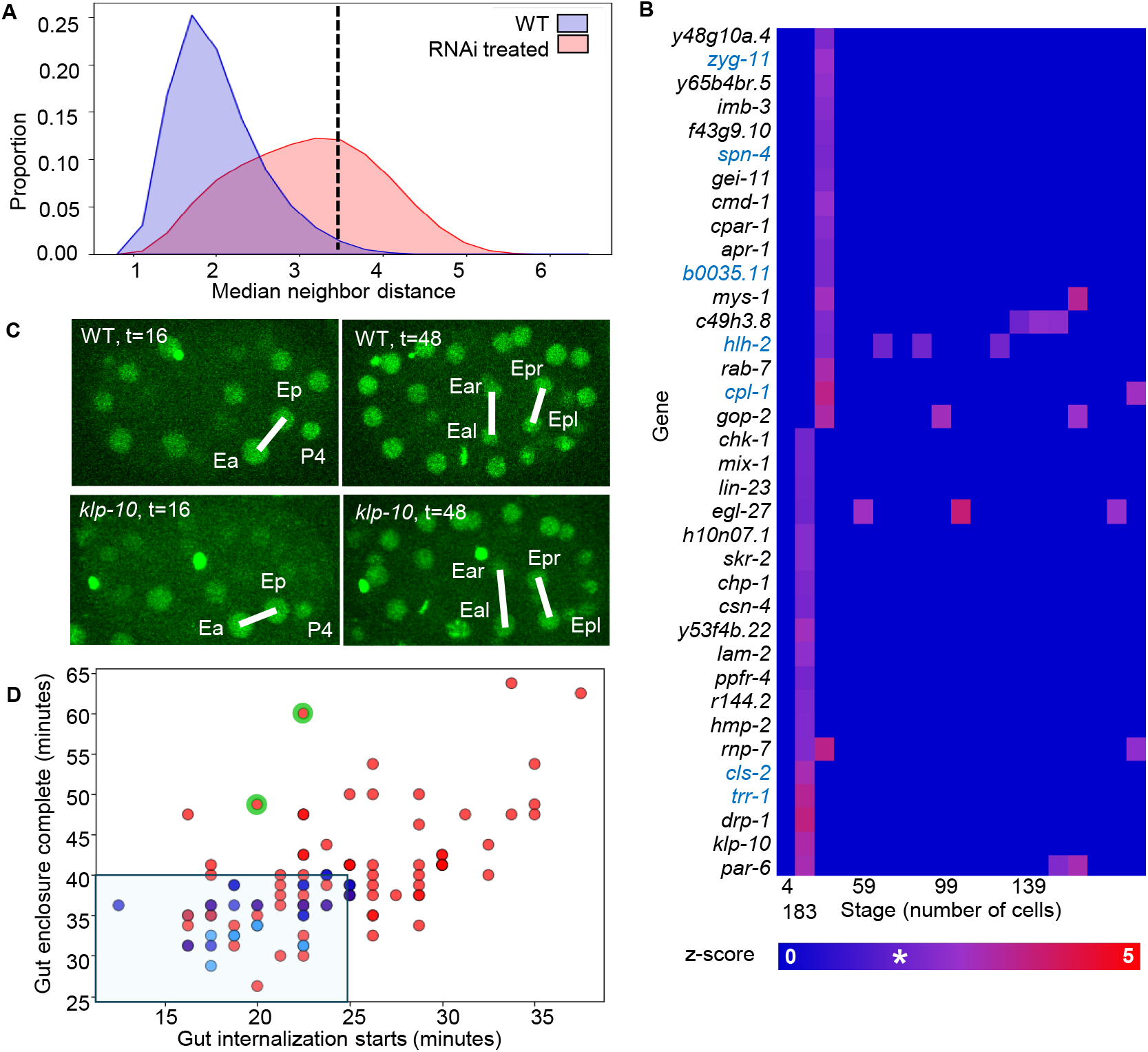
The developmental manifold enables complex morphology phenotyping from position data. A. Distribution of unusualness of an embedding measured by distance to the manifold (median distance to k=30 nearest neighbors). Manifold is of unedited WT embryos corresponding to dataset 1 (n=39). Distances are shown for all embeddings from an independent set of WT embryos (blue, n=10) and from embryos that survived weak RNAi treatment (red, n=703). Distributions are normalized to sum to one. A dashed line at 3.5 is chosen as a phenotype cutoff value. A value above this threshold indicates a phenotype. B. Gut phenotype heatmap. Intensity is a z-score for enrichment of phenotypes in predicted gut cells. The z-score compares the proportion of unusual gut cells in a location to the proportion of other unusual cells. This is binned by developmental stage (number of cells) across genes (n=144). Start and end bins are labeled with earliest and latest stage in the bin, respectively, and inner bins are labeled with the median stage per bin. Genes are sorted from bottom to top by stage of phenotype appearance. Only genes that are predicted hits for early gut development phenotypes are shown (see Figure S4.1B for all results). The * on the colorbar indicates the z-score associated with an alpha of 0.06, the 0.94 confidence level. Values below this cutoff are not shown. Displayed are the values for gene-representative embryos chosen as the embryo with the median (over the collapsed sum) severity per gene (n per gene varies from 1-23). Genes in blue text failed manual validation (Figure 5D). C. A maximum intensity projection of the middle third of a *klp-10* RNAi and a WT embryo. Times are in minutes post first gut lineage division. Key cells are labeled and sibling gut cells (based on lineaging) are connected by a white line. D. A scatter plot of gastrulation timing (Ep internalization (x) vs. enclosure (y)). See Figure S4.2A. Times are in minutes post first gut lineage division. Timings are shown for a set of WT (light blue) non gut hit RNAi (dark blue) and all hits from B (red). Two embryos are sampled per RNAi gene. *klp-10* data points circled in green.

To detect phenotypes, we use two levels of threshold: for individual cells, we use a data-driven threshold on distance from the WT manifold (3.5 median neighbor distances, Fig. 5a), and for lineage groups, a two-proportion z-score based on the proportion of cells over the first threshold (see Methods). Overall, phenotypes are detected in 132 of the 144 genes. The lineage groups are not affected evenly, ranging from 26 genes for the ABa group and 75 for MS (Fig. S4.1b), indicating lineage/tissue specificity for the genes. The majority of the genes also show a degree of specificity to developmental stages indicating transient phenotypes, which is consistent with the fact that the embryos hatched.

We focus on the E/gut lineage, which is known to undergo gastrulation, i.e., internalization, at this stage of embryogenesis. Confirmation of the results will not only validate our automated approach of phenotype detection, but also contribute novel regulators of gastrulation which have been difficult to find through screens due to genetic redundancy and lack of gross phenotypes [31, 32]. The automated analyses found 36 genes with E phenotypes at stages during and just surrounding gastrulation of the E cells (the 14-to 33-cell stages, Fig. 5b). We manually identify the E cells and quantify their timing of internalization (Fig. 5c and S4.2a). Comparing to the distribution in the WT and RNAi treated embryos where gastrulation was not affected (Fig. 5d and S4.2b,c), we found that for 29 of the 36 genes internalization was delayed (dots outside the shaded box in Fig. 5d) and for 7 (blue gene names in Fig. 5b), normal. Thus, the automated analysis has a true positive (TP) rate of 80% (29/36). These 29 genes show varying degrees of delay compared to the WT (Fig. S4.2d), from minor delays in initiation like *rab-7* and *apr-1* (Fig. S4.2e) to extreme delays in completion (not for another cell cycle) like *cmd-1* and *klp-10* (Fig. S4.2f). In all embryos the E cells ultimately gastrulate, as expected given embryos ultimately hatched. Function-wise, the 29 genes include cell polarity (*par-6*), cytoskeleton and regulators (*klp-10, cmd-1*), Wnt signaling or adherens junction (*apr-1, hmp-2*), protein degradation (*csn-4, lin-23, skr-2*) and others. Some of the genes are known or can be expected to affect gastrulation from the literature, eg, *par-6, apr-1* and *hmp-2*. Others may reveal new pathways regulating gastrulation upon further studies.

More broadly, extrapolating the 80% TP rate, our analyses suggest that about 100, or 2/3 of the genes in the dataset have tissue-specific transient morphogenesis phenotypes. In all, these results show that our approach localizes subtle and transient phenotypes in space and time despite the anticipated challenges, allowing automated screens and systematic discoveries on the regulatory mechanisms of tissue morphogenesis.

## Discussion

Our study successfully builds a learned spatiotemporal manifold that provides a unified and interpretable representation of single-cell tissue morphology and dynamics. Individual cells form distinct clusters. Spatially, the manifold preserves local distances, suggesting spatial configuration of neighborhoods and cell-cell interactions can be readily examined in the embedding space. Interestingly, embedding differences appear to be invariant to cell density in the real space, suggesting normalization to nearest neighbor distance as a strategy for unified measurement of morphology. This property, combined with the preservation of local distances, is reminiscent of a neighbor graph. However, the embedding also attends to global extrema, reminiscent of tissue axes. Temporally, the manifold is largely continuous describing a pseudo-time axis that is consistent with real time. Error types from cell type assignment in the embedding space, which are largely confusions between spatial or temporal neighbors, are consistent with these spatial and temporal properties of the manifold. Taken together, the manifold exhibits the desirable properties of a general descriptor, or universal language, to describe tissue morphology and its dynamic changes at single-cell resolution.

The applications further demonstrate the practical power of the manifold in interpreting tissue morphology and dynamic changes. As reflected along the way, the learned features are focused on appropriate structural features of tissue rather than over-specialized for training data, and effectively captures invariant relationships in highly variable data. The resulting model is richly descriptive, robust to data errors, and readily retrainable to create a foundational model, or for specific tasks to support experimental and mechanistic investigations.

Our success argues for a cell-based learning strategy as a productive and widely applicable alternative to the common image/pixel-based learning for building multi-modal foundational models [33, 34]. Logically, cells are the fundamental unit of tissue in which multi-modal data converge. Technically, cell point cloud-based manifolds offer the common spatiotemporal structure present across modalities, which is cumbersome to achieve in pixel-based learning. In practice, cell point clouds are readily available in live imaging of embryos, tissues and organoids, as well as single-cell spatial omics data. The matching task in our training approach can be generalized to learn on these samples. These samples often have many-to-many correspondences where each cell type has a variable number of cells. Set matching methods, unsupervised or weakly supervised, can solve the problem [35], and markers and omics data can be used to define equivalence classes of cell types. As more spatial omics data becomes available, this strategy can facilitate the correlative analysis of real-world imperfect patchworks of multimodal data sets. Finally, in terms of capacity to represent highly complex tissues, we note that in, seemingly simple, *C. elegans* embryogenesis over one thousand cell types through development are covered in our experiments, with hundreds present in any individual configuration and extreme variability, comparable to or surpassing the known complexity of typical tissue samples.

The Twin Attention approach may provide a powerful and general contrastive learning scheme beyond point clouds or tissue morphology. We postulate that part of our success in training a Transformer with limited data, both in quantity and breadth of sampling, is facilitated by the inter-sample attention more efficiently extracting variability, in contrast to typical contrastive loss schemes such as Siamese networks [36]. More broadly, the sampling aspect of Twin Attention essentially treats the data as N+1 dimensional; in our experiments N=3 (xyz) and time is the special dimension along which pairs are sampled. Generalizing the scheme, a single 3D volume can be sampled as pairs of 2D slices along the third axis. More broadly, dimensions of other natures across individuals such as experimental condition (e.g. concentration gradient, temperature), genotype or even species can all be used as the sampling dimension.

Our study is inspired by the success of molecular manifolds in organizing information about cell state and a desire to organize dynamic tissue morphology into a similar single-cell manifold representation. We believe our learned manifold provides a foundational representation for studying tissue biology, to synergize with molecular manifolds and other data modalities, and to systematically catalogue, compare and understand the patterns of cell relationships that structure tissue.

## Supporting information

Supplemental File 1 genes

## Acknowledgements

We thank Zhuo Du for initial data collection, Andrew Leifer, Christopher Brittin and Evan Vietorisz for valuable discussions and Murray Tipping for support. This work was partly supported by an NIH grant to ZB (R01GM152927) and an NIH Core Grant to MSKCC (P30 CA008748). AS was supported by Chan Zuckerberg Initiative Imaging Scientist Grant (2019-198110 (5022)).

## Author Contributions

Designed computational experiments: EH, AS, DW, ZB. Performed computational experiments: EH, AS, YX, RR. Performed imaging experiments and manual curation: YX, AS. Prepared manuscript: AS, ZB (with assistance from all authors). Supervised research: ZB.

## Software Availability

Training and inference code as well as pretrained models are available on Github: https://github.com/zhirongbaolab/TwinAttention/tree/main

## Data Availability

Edited and unedited cell position information for *C. elegans* embryos used in training and evaluating main results is deposited on zenodo: https://doi.org/10.5281/zenodo.16878148

## Methods

### Imaging and Lineage Analysis

All imaging data was collected as part of previously published studies [1, 25, 28]. All embryos were automatically segmented and tracked with the lineaging pipeline StarryNite as previously described [29, 30]. Manually curated datasets were edited in Acetree [37, 38] to establish correct lineage topology (assuring no tracks unexpectedly end, cross over, or unaccounted for cells appear) and hence establishing ground truth cell identity via automated naming rules.

### Simulation

An agent-based simulator [22] was modified to automate multiple simulation runs. The simulator utilizes data driven distributions of cell cycle timing to compute the probability of a division, apply average migration paths over a cell’s lifetime and use physical modeling to avoid collisions resulting in simulated nuclear centroids at one-minute intervals. 1000 simulations up to the 194-cell stage were generated with default parameter values.

### Preprocessing

Point clouds for each embryo are zero centered (by subtracting the mean of all points over all time) and converted from pixels to isotropic micron coordinates by multiplying each dimension by pixel dimensions. Each point cloud is then augmented by 10x with random 3D rotations around its centroid, to encourage rotational invariance.

### Sampling of Point Cloud Pairs

The sampling of ‘twin’ point clouds occurs across both individual embryos and over time. Developmental stage is established by number of cells. An adaptive sliding time window defines a series of overlapping equivalent periods within each embryo (Fig. 1b). Training pairs are randomly sampled from the union of the equivalent windows in each embryo and processed. The anchor in time of this window slides over development yielding a sampling of point clouds from different stages as it moves.

Training pairs are sampled from the data to cover all developmental stages while emphasizing the more challenging, rarer and more variable stages during cell divisions by over sampling them relative to stable developmental stages. We center an adaptive sliding window *W* at each stage *i* (defined by number of cells in the embryo) and draw *S* samples from the point clouds augmented by rotation available in this window. We define *S_sufficient* data for a stage empirically as 3*number of training embryos*number of random rotations. Number of embryos=33 for real data (experiment 1) 155 for real data (experiment 2) and 1000 for simulated data. Samples drawn *S*=min(S_sufficient, samples_in_window). Samples are randomly divided into batches of 10 and at each epoch these batches of samples are turned into batches of training pairs by shuffling, choosing the first set in a batch and concatenating it with each other set as a comparison reference embryo.

The adaptive window size *W* from which these samples are drawn is a function of the number of samples at and surrounding the anchor stage *i. W*=1 if samples(*i*) > S_sufficient. This indicates the stage is a developmental plateau that is already densely sampled. Otherwise *W* is grown till it hits a max size or an adjacent ‘sufficient’ plateau stage. *W* is independently expanded by one in each direction until an adjacent stage in that direction has samples >=S_sufficent or the window attains a maximal radius.

Window max radius (*j*) is stage dependent: stage<=50 cells: *j*=2, stage>50<=100 cells: *j* =5, stage>100<=200 cells: *j* =10, stage >200<=300 cells: *j* =15,stage >300 cells: *j* =25. This reflects roughly the decreasing importance of a single-cell increase in stage as the total number of cells increases.

Training Data Summary.

Training Data Summary
| Dataset | Training | Training Test | Held-Out |
| --- | --- | --- | --- |
| Simulations | 1000 | 100 | 10 |
| Dataset 1 | 33 | 6 | 10 |
| Dataset 2 | 155 | 29 | 49 |

### Model Architecture and Training Hyper Parameters

Our model consists of an MLP which up samples 3D positions to a 128 dimensional embedding, followed by a standard Transformer Encoder layer implemented on top of [11]. As in [11] the two sets are distinguished when concatenated by a uniform offset, akin to a positional encoding, added into one point set. Loss followed the previously published scheme, essentially cross entropy loss on the softmax of the cosine distance of embeddings of each set and the correct matching. For training, daughter cells are treated as having a distinct identity from their mother cell. The training loop starts with a base learning rate and decreases learning by a multiplicative factor of learning rate decay whenever patience batches have been processed without a decrease of loss. After max trials reductions in learning rate training stops early.

| Parameter | Setting |
| --- | --- |
| Embedding dimensionality | 128 |
| Transformer layers | 6 |
| Transformer heads | 8 |
| Batch size | 10 |
| Learning rate decay | .25 |
| Base learning rate | $2e^{-4}$ |
| Patience | 10 |
| Max trials | 20 |

### Inference

To create embeddings to represent an embryo post training we generate a batch of size one by concatenating the query embryo with a random stage matched embryo from the *training test* portion of the relevant dataset. This pair is processed. For pairwise matching both pairs are used, with identities from training dataset attached to the matched cells in the query data. When generating the developmental manifold, embeddings for the query are retained while the reference is discarded. Named reference developmental manifolds for all subsequent analysis were generated using all training data point clouds (train and train test).

### Ablation

In ablation experiments a cell’s entry in the input 3D position matrix is completely removed and the reduced set of points is embedded. Embeddings for remaining cells pre and post ablation are compared.

### Pseudo-time Analysis

Embeddings for a cell over generations were gathered from three embryos. These embeddings were processed with a diffusion map algorithm (pydiffmap, epsilon=‘bgh’ k=#points/2. Ordering or position along the resulting manifold were used as pseudo-time in respective plots.

### Cell Type Assignment

While it is possible to assign cell types by matching with a single labeled template, uncertainty always exists whether a given template will produce optimal results. This problem is exacerbated in development where heterochrony means different cells can be present in similar templates (see Fig.S1c,d). Doing multiple alignments and voting [11, 19] or optimizing template contents [19] mitigates, but does not remove dependence on template choice. KNN classification using the reference manifold is the simplest way to use all available information simultaneously.

The full set of reference manifold embeddings over all stages were used as an input distribution to a KNN classifier with cell identities as class labels and k=30 (optimized with exhaustive testing). The fine tuned model for dataset 1 was used in cell type assignment (for dataset 1) and subsequent phenotyping experiments as its training data was a closer match to the imaging conditions and developmental stage of the RNAi data.

### Phenotyping

For phenotyping of uncurated embryo movies an unedited reference WT manifold was generated by embedding the *unedited versions* of the WT embryos used to create the cell assignment manifold. Cell identities are propagated from the edited to unedited point clouds by nearest neighbor matching in real 3D space between individual edited and unedited point clouds. The encoding of unedited WT embryos creates a more accurate view of how WT behavior will look when corrupted with occasional detection errors. This also allows the creation of an ‘other’ identity (to be ignored) that can be attached to cells in the WT manifold (cells not corresponding to a TP in the ground truth): origins of these cells include detection errors, polar bodies, or cells from adjacent embryos in the field of view. This helps prevent these artifacts in mutant embryos from being identified as some other improbable, but closest cell type (as would occur if they were named against the *edited* WT manifold).

Since multiple embryos are present in a field of view, cells from other embryos sometimes add distant detections to unedited results (analysis is limited to a bounding box which sometimes overlaps with other embryos). To minimize these all cells at all timepoints were clustered with the dbscan algorithm (epsilon=3) in MATLAB. Only the largest cluster (corresponding to the complete embryo that was the intended focus) is retained. This procedure was followed for unedited WT and RNAi embryos.

All unedited RNAi embryos were embedded, their names predicted using the WT distribution and the median of their k=30 NN recorded. To define unusual embeddings distance was calibrated to a threshold of 3.5 median neighbor distance. Past this there are a negligible number of embeddings from the WT distribution. The count of embeddings at each stage (binned at ten cell intervals) and for each founder lineage (cells descended from a major progenitor, here specifically the set of embryonic progenitors ABa, ABp, E, MS, P2) in each embryo was tallied. These counts were converted into a z-score by computing a two-proportion z-score for each stage and lineage location. A particular time/founder lineage location is designated as L and a z is computed for the difference between ratios: unusual(L)/unusual(not L) and usual(L)/usual(not L). This captures the intuition that the prevalence of unusual embeddings in that location is above the background prevalence of that cell type within the embryo. For each gene an ‘exemplar’ embryo is picked for visualization, as the embryo with median total unusual embeddings over all lineages. This avoids the risks averaging over distinct phenotypes for a RNAi condition due to penetrance variation.

To identify potential gastrulation phenotypes for validation genes with z-scores above.06 in either the second or third binned time stage were selected, that is, the 14-33 cell stages encompassing the onset of gastrulation which in the WT is at the 24-cell stage.

### Phenotype Validation

To validate gastrulation phenotypes embryos from several conditions were measured, 2 randomly selected embryos were measured of each RNAi hit gene as well as 10 WT embryos and 2 embryos each of 10 randomly selected non-hit RNAi genes.

In each of these embryos the E gut cell was manually identified and followed. The time point before E initially divides defines time zero. The two E cells were manually followed forward and the time of Ep completing its ingression was recorded. As seen in Fig. S4.2a this stage is characterized by Ep having entered while Ea remains on the surface, maximizing the angle between the two and P4. In the WT this results in a distinctive 90 degree angle between Ea,Ep and P4 (identified by its position at the posterior end of the embryo). In RNAi embryos this angle is not always as sharp but the moment with maximal angle between the three cells is taken (later, this angle is reduced by Ea internalizing). The E cells were followed forward till the ingression of Ea and Ep were complete. As it is sometimes difficult to assess the motion of the gut cells once they are no longer on the surface the completion of gastrulation was defined as when surrounding cells had covered the ingressed E cells with P4 and MSap having moved to fill the space left by Ea/p. This event is recorded as the first frame after both E cells are off the surface in which the angle Ea-MSap Ep-P4 stops decreasing, with an angle of 30 degrees typically remaining in the WT.

---

**Figure S1.**
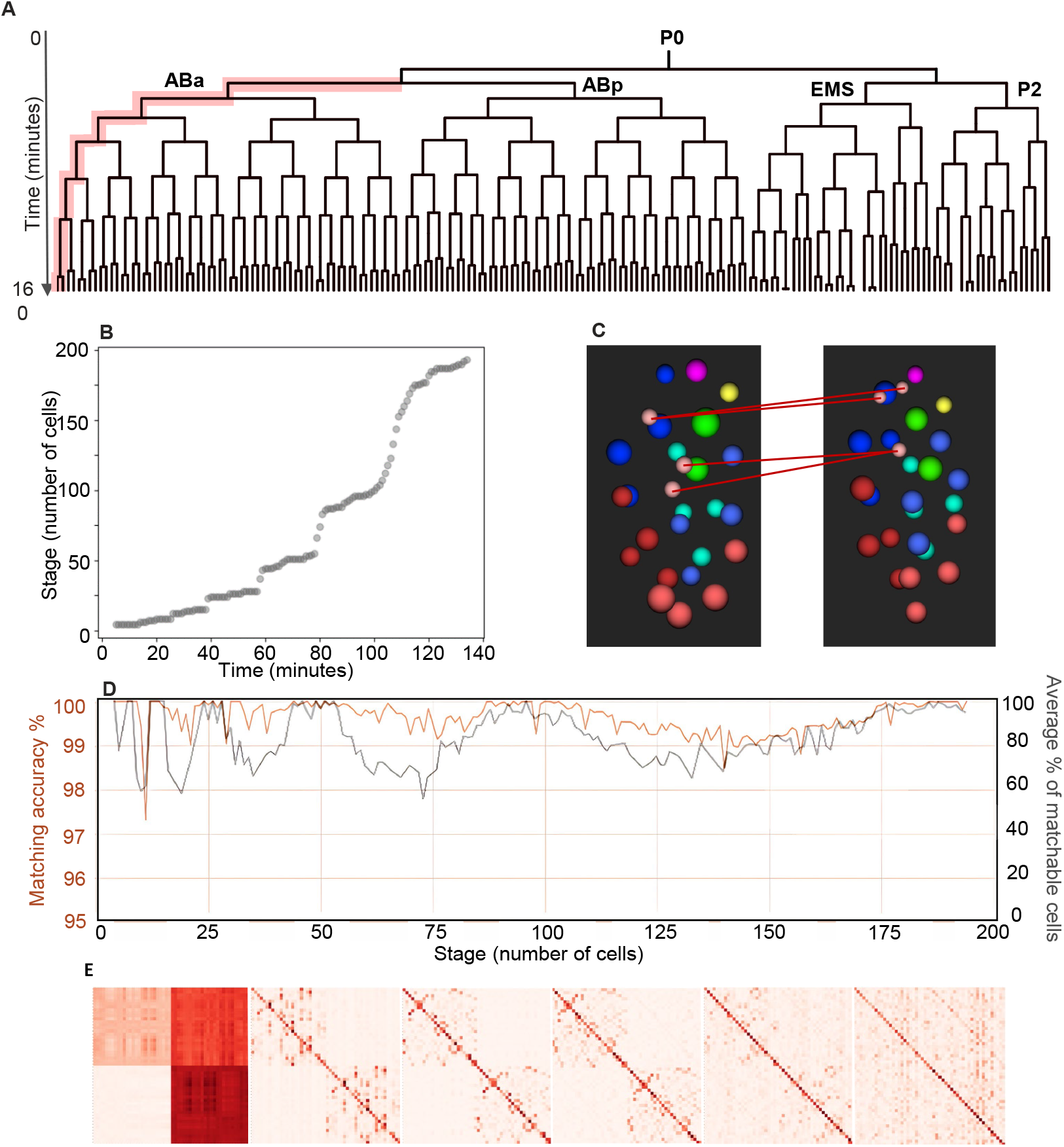
Learning a single-cell developmental manifold of tissue morphogenesis, continued. A. The early *C. elegans* cell lineage. Vertical lines represent cells with length proportional to time. Bifurcations represent cell divisions. Founder cell names are labeled, with P0 being the zygote. Red highlights the concept of lineage path by labeling an example lineage path (ABa-ABalaaaa). B. Time versus the number of cells present in a typical developing embryo. C. Heterochrony. A pair of embryos around the 26-cell stage. Divisions of the C lineage are labeled with red lines connecting mother cells in one embryo to corresponding daughters in the other. D. Matching accuracy on holdout data (orange), calculated for the matchable cells between a pair of embryonic point clouds, and the average ratio of matchable cells between pairs of embryonic point clouds across developmental stages (gray) (n=25 random sample pairs per stage from 10 held-out embryos). E. Attention maps computed at each of the 6 layers of the Transformer encoder between ***E***_**1**_ and ***E***_**2**_ as in Figure 1D. Maps are ordered from left to right by increasing depth in the model.

**Figure S2.**
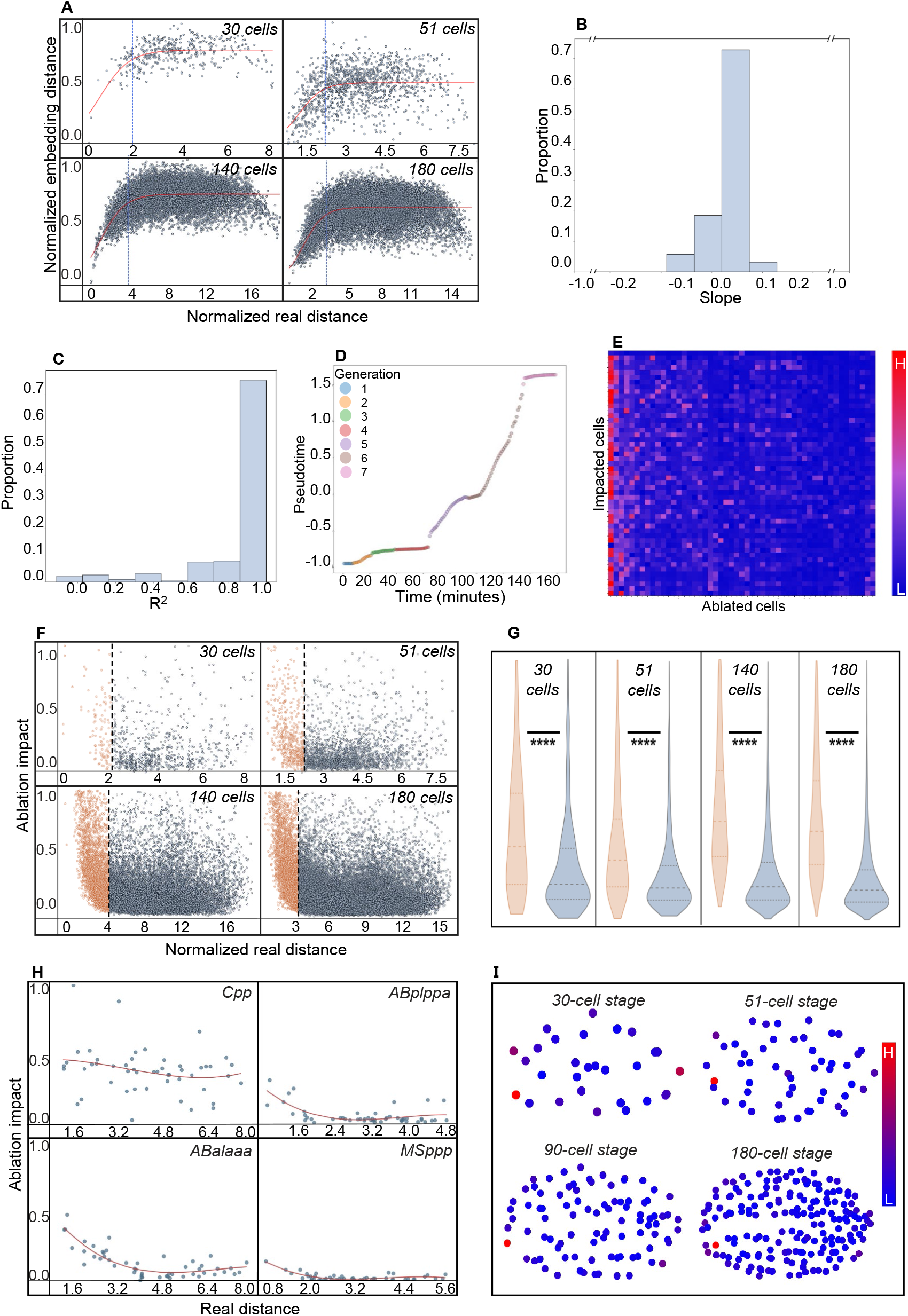
Characterization of the developmental manifold, continued. A. Embedding distance reflects local distance in real space. Pairwise cell distance in real 3D and embedding space. Shown are 30,51,140,180 cell-stage embryos. Red line is a fit with a sigmoid function. Normalization as in Figure 3A. B. Scaling with density. Histogram of slopes for a linear fit of distance between siblings in embedding space over generations as in Figure 3B but computed along every lineage path from the 4-194-cell stage (n=1 embryo). C. A histogram of the R^2^ values for a linear fits of the relationship between time and pseudotime as described in Figure 3C (every lineage path from the 4-194 cell stage (n=1 embryo)). D. Pseudotime axis. Time versus 1D diffusion map coordinates of cell embedding over the AB-ABalaapaa lineage path (colored by cell and numbered by generation, n=1). E. In silico cell ablation impact. A heatmap of computational cell ablation results (‘ablation impact’). Column indicates ablated cell. Results are shown for the 51-cell stage, with columns sorted by hierarchical clustering. F. Local influence of in silico cell ablations. Influence of cells at varying distances on embeddings. Pairwise cell distance in real space versus the change in embedding position over multiple stages as in Figure 3D. G. A summary of ablation impact in (F), formatted as in Figure 3E. H. Strong local influence of in silico cell ablations is consistent across cells. Pairwise real distance vs ablation impact on a single cell from ablation of all other cells. Third-degree polynomial fit in red. Real distance is normalized by the average nearest-neighbor distance across all cells at the stage. I. Some cells show global impact. Summary of per cell ablation impact shown at different stages as in Figure 3F.

**Figure S3.**
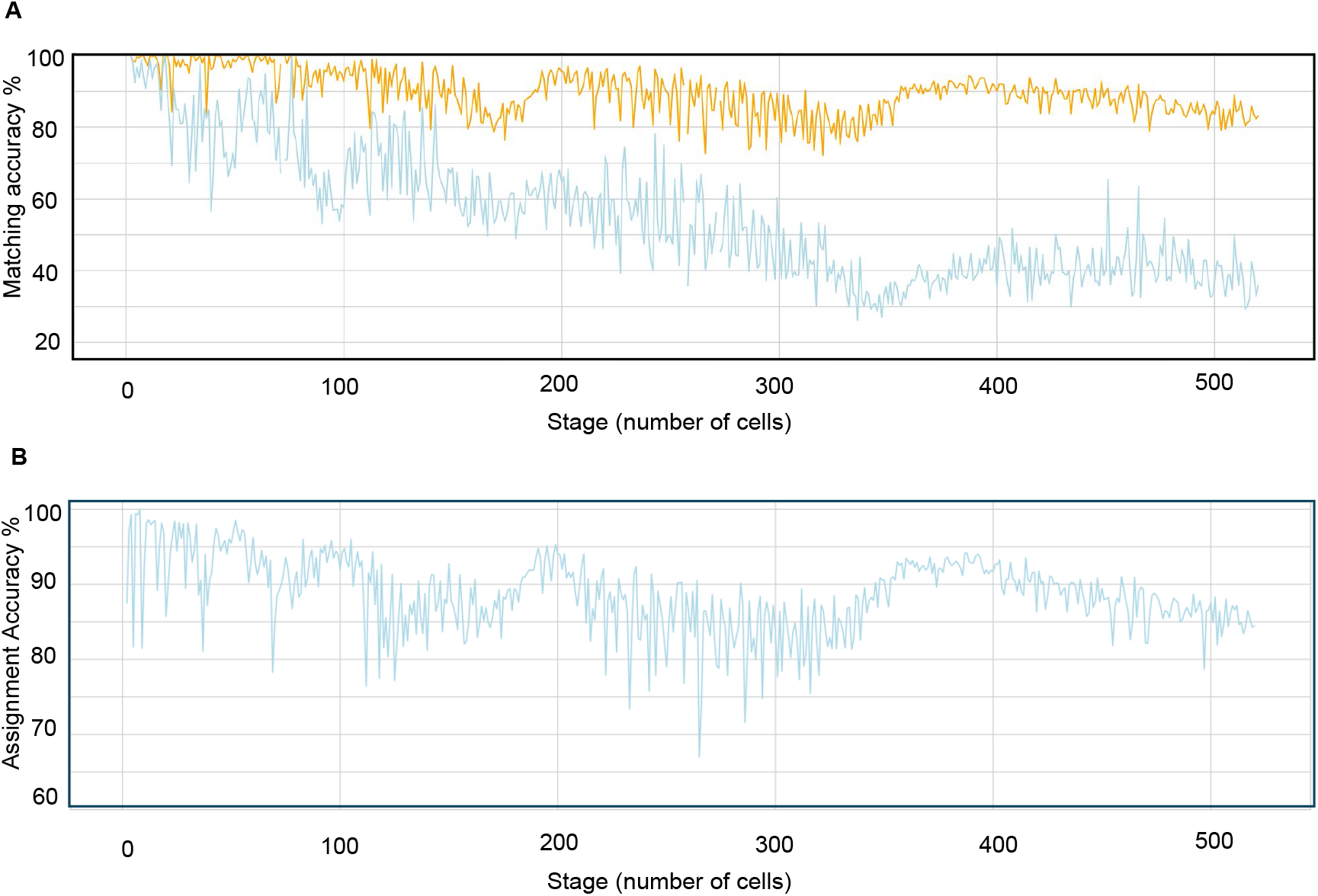
Fine-tuning of base model for cell type assignment, continued. A. Pairwise matching accuracy for matchable cells over stages on experimental dataset 2. Results are shown for the model fine-tuned using experimental dataset 1 (blue) and experimental dataset 2 (orange), while testing in both cases uses the held-out portion of experimental dataset 2 (n=200 random sample pairs from 49 held-out embryos for each stage). B. Matching accuracy from kNN classification over stages, experimental dataset two (n=49 held-out embryos)

**Figure S4.1.**
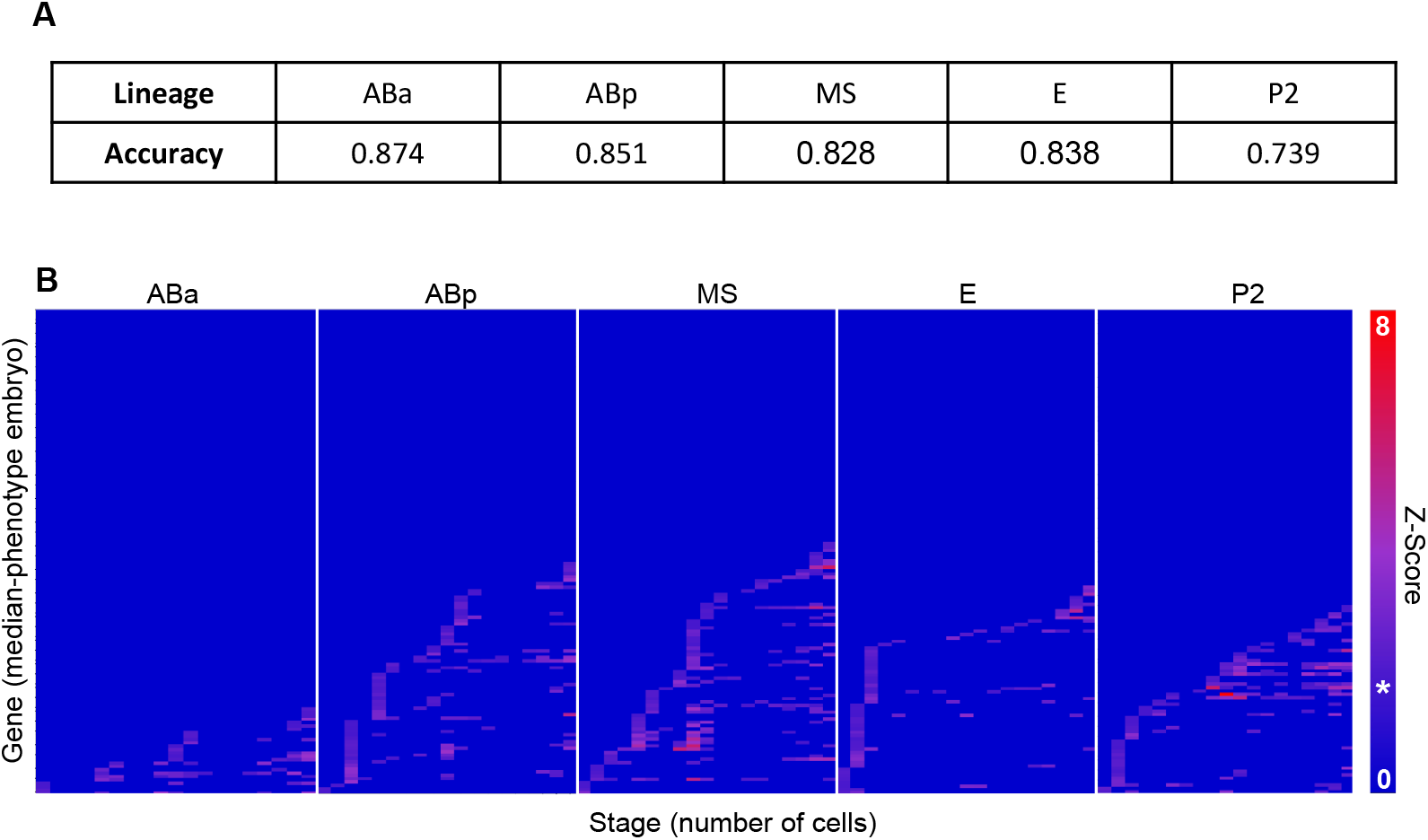
The developmental manifold enables complex morphology phenotyping from position data, continued. A. Lineage-level assignment accuracies (TP rate, correct/all predicted for that lineage) in the subset of RNAi treated embryos with manually validated identities (n=32 embryos 4-194 cell stage). B. Phenotype heatmaps for major embryonic sublineages. Color indicates z-score for enrichment of unusual embeddings for each predicted founder lineage, developmental stage (number of cells) and gene (n=144). Genes are sorted (independently for each founder) from bottom to top by stage of phenotype appearance. The * on the colorbar indicates the z-score associated with an alpha of 0.06, the 0.94 confidence level. Values below this cutoff are not show. Gene-representative embryos shown have the median (over the collapsed sum) severity out of all embryos for that gene (n per gene varies from 1-23).

**Figure S4.2.**
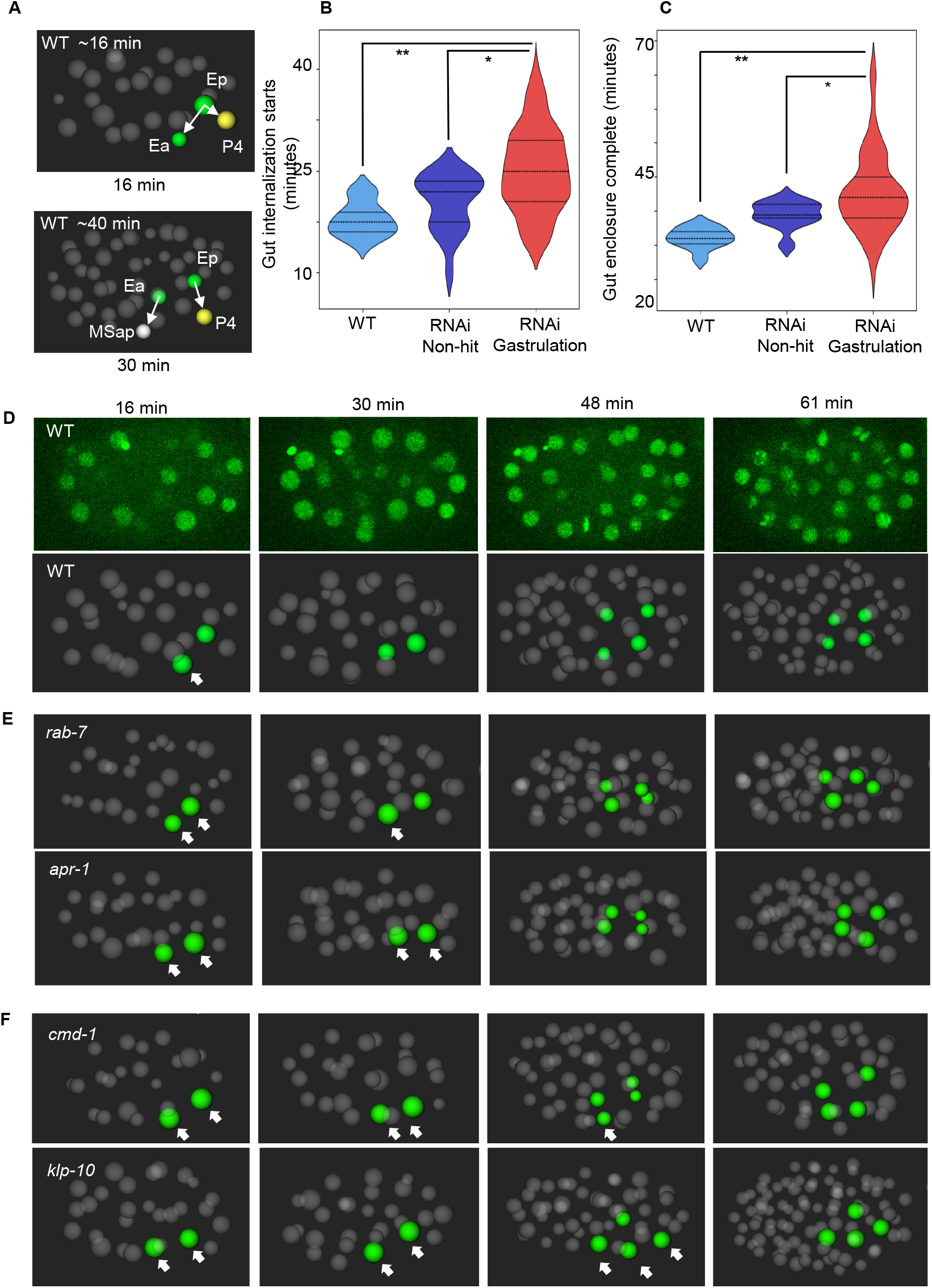
Embeddings enable complex morphology phenotyping from position data, continued. A. Illustration of configurations timed in gastrulation phenotype validation. First landmark: Ep internalizes, forming a maximal angle with Ea and P4 (∼90 degree in WT) before Ea’s internalization reduces the angle. Second landmark: both E cells are internalized and P4 and MSap have moved to cover them. This is defined as the earliest time at which lines Ea-MSap and Ep-P4 define a minimal angle to each other (∼30 degrees in WT). B. Timing of Ep internalization (minutes after first gut division) in WT (light blue, n=10), RNAi treated embryos negative for a gastrulation phenotype (dark blue, n=20), and RNAi embryos positive for gastrulation phenotype (red, n=71). The gastrulation hit RNAi treated embryos are significantly delayed compared to WT (**, p=0.0005) and the non-hit RNAi treated embryos (*, p=0.002). C. Time of full gut precursor enclosure (minutes after E division) in WT (light blue, n=5), RNAi-treated embryos negative for any gastrulation phenotype (dark blue, n=10), and RNAi embryos positive for gastrulation phenotype (red, n=71). The gastrulation hit RNAi treated embryos are significantly delayed compared to WT (**, p=0.0006) and the non-hit RNAi treated embryos (*, p=0.02). D. Progression of gastrulation in a WT embryo (as in Fig. 5c) Shown as a maximum projection of the middle third of the embryo and as a 3D rendering from the same dorsal direction (gut precursors in green, exposed gut cells marked with white arrow). E. Progression of gastrulation in two RNAi embryos (at same times post first gut cell division as WT in D). These show shorter delays where gastrulation is complete before an additional round of gut precursor divisions (gut precursors in green, exposed gut cells marked with white arrow). F. Progression of gastrulation in two RNAi embryos with more severe delay in which one or more of four gut cell precursors are born on the embryo exterior.

